# Accumulation of dead cells from contact killing facilitates coexistence in bacterial biofilms

**DOI:** 10.1101/2020.04.03.024158

**Authors:** Gabi Steinbach, Cristian Crisan, Siu Lung Ng, Brian Hammer, Peter Yunker

## Abstract

Bacterial communities govern their composition using a wide variety of social interactions, some of which are antagonistic. Many antagonistic mechanisms, such as the Type VI Secretion System (T6SS), require killer cells to directly contact target cells. The T6SS is hypothesized to be a highly potent weapon, capable of facilitating the invasion and defense of bacterial populations. However, we find that the efficacy of the T6SS is severely limited by the material consequences of cell death. Through experiments with *Vibrio cholerae* strains that kill via the T6SS, we show that dead cell debris quickly accumulates at the interface that forms between competing strains, preventing contact and thus preventing killing. While previous experiments have shown that T6SS killing can reduce a population of target cells by as much as one-million-fold, we find that as a result of the formation of dead cell debris barriers, the impact of T6SS killing depends sensitively on the initial concentrations of killer and target cells. Therefore, while the T6SS provides defense against contacting competitors on a single cell level, it is incapable of facilitating invasion or the elimination of competitors on a community level.

## Introduction

Bacteria commonly inhabit biofilms in the form of crowded, surface-attached microbial consortia embedded within a viscous matrix of polymers. Interactions between different bacterial strains and species govern the spatial organization and composition of biofilms [1–3], and ultimately affect the proliferation and survival of individual strains. These interactions can turn deadly. Bacteria have evolved many mechanisms to kill each other within biofilms [4, 5]. Several of these killing mechanisms require direct contact between cells, which typically occurs only within crowded biofilms [4, 6–9]. One such contact killing mechanism is the broadly prevalent Type VI Secretion System (T6SS) in Gram-negative bacteria [10]. A significant amount of work has produced a detailed picture of the T6SS. Details are emerging of the T6SS structure, toxins, and regulation [10–20]. However, the importance of this lethal activity in natural communities remains unclear [21, 22]. Experiments have primarily focused on the outcome of competitions between T6SS-proficient (T6SS+) ‘killers’ and target strains that lack T6SS activity, but the dynamics of T6SS killing are much less studied [23] (though dynamic simulations have made a number of successful predictions [24–28]). Understanding the impact of the T6SS requires experimental observation of contact killing in microbial communities as a function of time and isolated from other factors. This is a crucial step in assessing the ecological role of T6SS-mediated contact killing over short and long time scales.

The T6SS is widely considered a potent weapon. In previously described killing competition assays, T6SS-proficient bacteria decrease the abundance of target cells by as much as one-million-fold within a few hours [15, 23, 29–31]. Based on these results, the T6SS is hypothesized to play important roles in inter- and intra-strain competition, for example, facilitating invasion of colonized space, elimination of competitors, and defense against invaders and cheaters [21, 28, 32–34]. However, competition assays also suggest that T6SS-mediated killing is rarely able to completely eliminate all target cells, despite drastically reducing their abundance. Even when killer cells start at numerical advantage (a 10:1 number ratio of killer:target cells is often used for competition assays), some target cells nearly always survive [23, 24, 30, 31, 35]. Further, though the killing rate is typically very high at the start of an experiment, killing almost halts after a few hours, despite the presence of target cells; this dramatic decrease in killing occurs even when the killer strains expresses a constitutively active T6SS [23]. It is difficult to isolate alterations in T6SS activity over time as developing biofilms become increasingly heterogeneous [36, 37] and constantly change [38–42]: biomass increases, nutrient and oxygen concentrations drop, excreted waste products accumulate, cellular behavior changes due to signaling from secreted public goods. As a result, a detailed picture of how T6SS killing proceeds within biofilms, and why T6SS active strains do not completely eliminate their competitors, remains elusive.

Here we show that while T6SS-mediated killing is very effective on first contact between competing strains, killing nearly ceases after a few hours due to the accumulation of dead cell debris. Contact killing experiments typically focus on living cells as they measure the number of surviving target cells [15, 29–31, 43–47]. This approach was fundamental in discovering the deadly effect of T6SS [7], and the characteristic spatial structure that emerges from contact killing [2, 24, 26, 28]. Here, we focus on the role of dead cells and dead cell debris. Through microscopy experiments with *V. cholerae* we confirm that dead cell debris accumulates at the interface between the clonal domains of competing strains, which form upon killing and reproduction [24]. We find that dead cell debris eventually prevents contact and thus halts killing. Paradoxically, contact killing may thus play a protective role in biofilms, facilitating the formation and coexistence of separate clonal domains [48, 49].

## Results

To study how T6SS killing proceeds within biofilms, we optically record the spatio-temporal dynamics of biofilms comprising two engineered strains of *V. cholerae*, a model organism for studying the T6SS [7]. These strains are isogenic, and only differ in their T6SS toxins and immunity modules, their ability to express T6SS, and their fluorescent proteins (see methods) [24]. To isolate the effects of killing from other effects related to changes in biofilm height (which can impact cellular behavior [37]), we grow biofilms in confinement between an agar pad and a cover glass (Fig.1a) (see methods).

As a first step, using confocal time-lapse microscopy we investigate the temporal dynamics of unidirectional killing: the killer is a T6SS-proficient (T6SS+) *V. cholerae* strain, while the target is a susceptible, T6SS-defective (T6SS-) *V. cholerae* strain. We observe the formation of clonal domains (Fig. 1b) upon reproduction and killing. This phenomenon is reminiscent of domain formation observed in populations of mutual killer strains (i.e, both strains are T6SS+) [24], even though in our current experiments one strain – the target strain – is engineered to be defective at T6SS killing. While we observe that domains of the T6SS+ strain expand quickly at early times, domain growth later slows substantially (Fig. 1b). We quantify the temporal dynamics of mixed populations of killer and target strains by measuring the relative abundance of the fluorescent killer strain in the microscopy images over 15 hours, for four different initial ratios of killer and target cells (Fig. 1c), see methods. In all cases, the killer strain initially increases its population rapidly; the killer population reaches ∼ 90% of its final size after ∼ 3 hours. Killing dramatically slows afterwards. We observe similar temporal dynamics in experiments with mutually killing strains (S2 Fig): again, a transition from rapid killing to almost no killing occurs after ∼3 hours (S1 Fig b). These observations are consistent with previously reported observations of small but long-lived target populations [23]. Why does T6SS-mediated killing stop after just a few hours?

**Fig 1.**
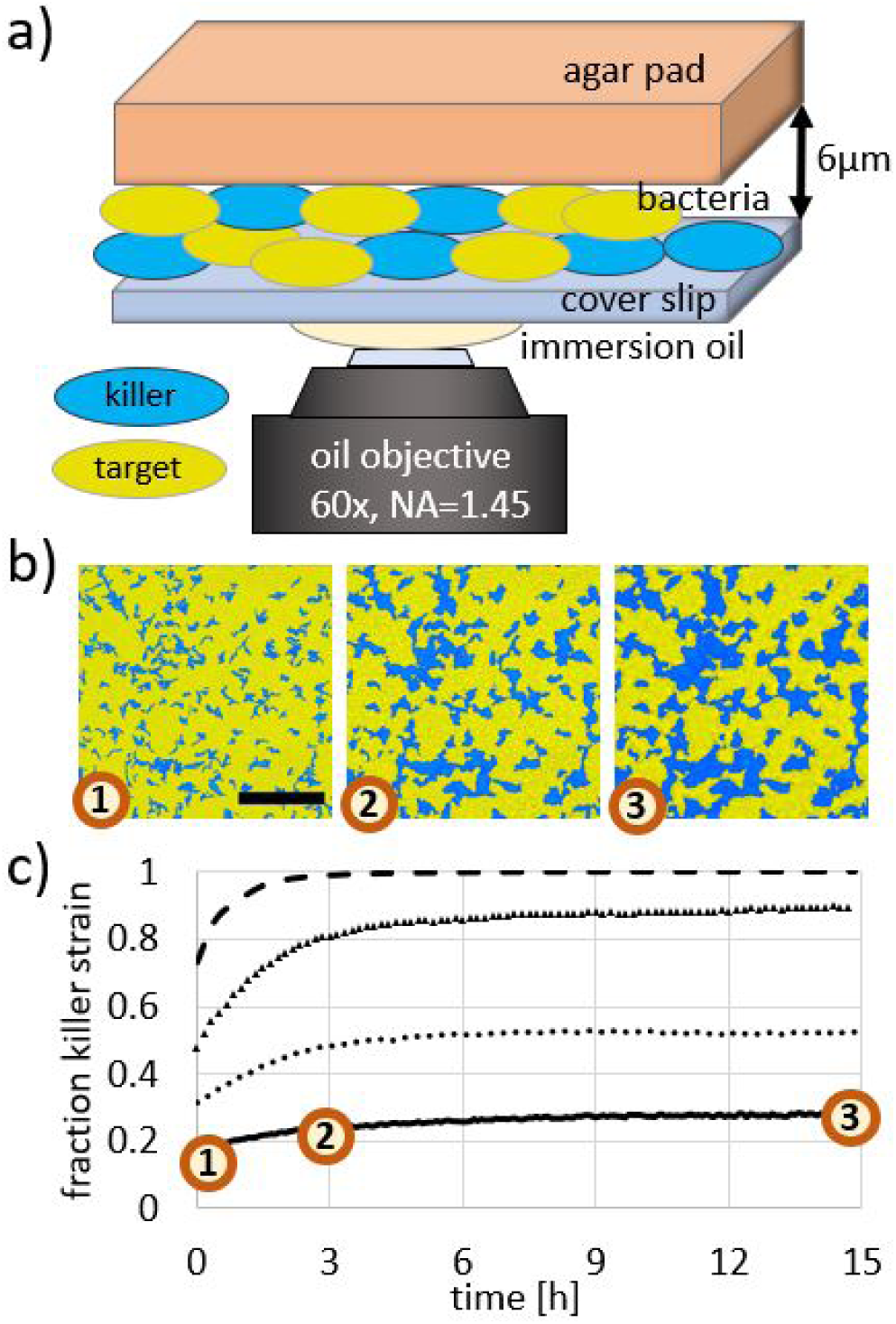
Contact killing slows substantially over time. a) Cartoon depiction of biofilm samples for imaging. Confinement between an agar pad and a glass cover slide leads to a maximum biofilm thickness of ∼6*µ*m. b) Fluorescence images of a unidirectional competition between killer cells (blue) and target cells (yellow) at 0, 3, and 15 hours (left to right, respectively). The initial relative abundances were 0.09 killer cells and 0.91 target cells. T6SS-mediated contact killing leads to the formation of clonal domains, which increase in size over time. Scale bar: 50*µ*m. c) Relative abundance of the killer cells over time for four different initial conditions (killer cell relative abundance during inoculation: 0.5, 0.33, 0.17, 0.09, from top to bottom curve) measured from time lapse images as depicted in (b). In all cases, the relative abundance of killer cells initially increases quickly, followed by a substantially slower increase after 3 hours.

Surprisingly, we find that the boundary between competing strains is visible in images recorded with bright-field microscopy (Fig. 2a). The interface between domains of killer and target cells in fluorescence images (Fig. 2a, bottom image) aligns with dark outlines that are visible in images taken in bright-field mode (Fig. 2a, top image). The ability to visualize the interface between competing strains using bright-field microscopy is not expected since the two isogenic strains don’t differ in their material properties—including index of refraction—so they appear identical when imaged with bright-field microscopy. In fact, such dark outlines are absent in biofilms without killing, i.e., those with two isogenic T6SS-strains. Small clonal domains still emerge as non-motile, divided cells typically remain close after reproduction (Fig. 2b, bottom image). However, the interface between T6SS-clonal domains does not appear dark in bright-field images (Fig. 2b, top image). The presence of dark lines in biofilms with T6SS killing suggests that there is a change in material properties (e.g., index of refraction) at the border between strains. Based on this observation, and the data presented above, we hypothesize that dead cell debris accumulates as cells are killed. Such cell debris may eventually prevent competing cells from contacting. Similar observations have been made previously when studying T6SS-mediated interactions [50–52].

**Fig 2.**
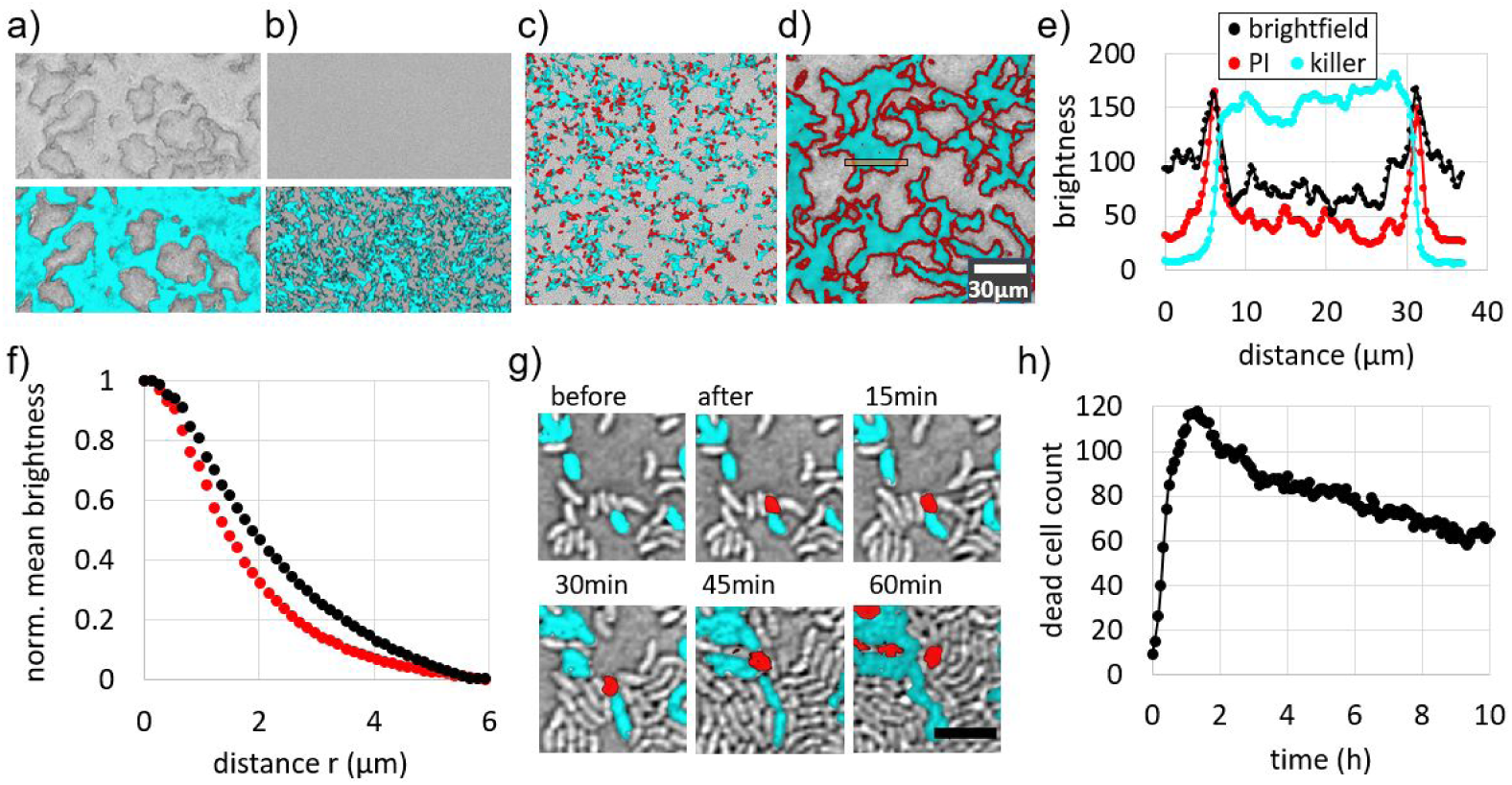
Characterization of dead cell debris. a) Representative bright-field image alone (top) and overlayed with fluorescence channel (bottom) of biofilms exhibiting unidirectional killing, recorded after 7 hours of growth. The killer expresses sfGFP (cyan) while the target is unlabelled. Note, the entire field of view is densely packed with bacteria. b) Same as (a) but the killer strain is engineered to be T6SS-deficient; no T6SS killing occurs here and dark outlines are absent. c, d) Merged bright-field and fluorescence images of unidirectional killing after 1 hour (c) and 7 hours (d) of growth. Again, the killer expresses sfGFP (visible as cyan) while the target is unlabeled (gray). The DNA of compromised cells, i.e., dead cell debris, is labelled red via PI. e) Intensity profile of different microscope images across a clonal patch (integrated vertically over the orange box in (d)) in the fluorescence images and bright-field image (inverted grey scale). f) Normalized mean intensity of PI signal (red curve) and inverted bright-field signal (black curve) within a distance *r* from the interface between competing strains in (d). g) Killing event: a fluorescent killer cell kills a non-fluorescent target cell, which subsequently turns red. The dead cell debris is relocated by neighboring cells that exert forces upon growth and reproduction. (Scale bar: 4*µ*m) h) Counting the number of PI labeled cells over time demonstrates that PI-labeled dead cell debris persists long after cell death. Data are recorded from individual target cells densely surrounded by killer cells (initial target to killer number ratio of 1:50).

To test our hypothesis, we track dead cell debris in growing biofilms with propidium iodide (PI), Fig. 2c,d (see also S3 Fig). PI binds to the DNA of cells with a compromised membrane and exhibits high red fluorescence. While stained dead cells appear throughout the biofilm during the earliest stages of growth when clonal domains are small (Fig. 2c), at later times the dead cell stain is clearly localized at the interface between large clonal domains (Fig. 2d). The PI signal is very well aligned with both the interface between strains and the dark outlines seen with bright-field microscopy (Fig. 2e). The PI signal exhibits a peak at the same position (distance of 6.2*µ*m and 31.3*µ*m) where the cell fluorescence declines to about 30% of its maximum value. From a Gaussian fit we find that both peaks in PI signal and bright-field signal differ in position by less than 0.3*µ*m and differ in width (i.e. standard deviation of the peak) by less than 0.1*µ*m. This sub-micron alignment of signals suggests that dark outlines observed via bright-field microscopy correspond to a substantial amount of cell debris at the interfaces between patches.

To quantify the localization of dead cell debris at interfaces throughout the biofilm, we measure the mean intensity of PI signal as a function of distance from the interface between strains (Fig. 2f, see methods for more details on the analysis). The intensity of the PI signal decays with distance from the strain interface, and reaches half its maximum value at a distance of 1.5*µ*m. We apply the same image analysis to bright field images and characterize the dark outlines at the strain interfaces. The dark outlines lead to a similar decaying curve, reaching half its maximum value at a distance of 1.8*µ*m. Both curves confirm that dead cell debris is highly localized at the interface between strains. Crucially, the average dead cell debris layer thickness of 1.5*µ*m is larger than the length of a *V. cholerae* cell.

To account for the observed slow rate of killing after three hours, the dead cell debris that separates competing strains must also be stable over long periods of time. We observe that debris from one individual dead cell can remain clearly visible for at least 60 minutes, even as it is relocated via forces exerted by neighboring cells as those reproduce and die (Fig. 2g). However, the emergence of more dead cells in close proximity inhibits tracking for longer times.

To quantify the stability of many dead cells over long times, we mix 98% T6SS+ killer cells and 2% T6SS-target cells and inoculate at high density (OD_600_ = 10), so experiments begin with close-packed cellular monolayers. Target cells are very far from each other, allowing us to isolate and track individual stained dead cells over long times (∼ 10 hours). All tracked cells (∼ 120) are killed within 1.5 hours of inoculation (Fig. 2h). Afterwards, the number of dead cells with detectable PI signal slowly decreases over time. The decrease in the number of PI-labeled dead cells may be due to a local loss in the presence of dead cell debris, e.g., the material may degrade and diffuse away. Note that the PI-labeled area per dead cell also decreases over time (S2 Fig), which may be caused by degradation or compaction of dead cell debris. However, despite the observed decrease of dead cell debris, over 80% of dead cells display a clear PI signal after 4 hours, and the majority of dead cells (over 50%) still display a clear PI signal after 10 hours. Thus, a substantial amount of dead cell debris persists over long times.

Accumulation of dead cell debris is also present in competitions between non-isogenic strains. To test the accumulation of debris between non-isogenic strains, we compete the previously used *V. cholerae* killer (T6SS+) against other killer strains (four other environmental isolates of *V. cholerae* [29]) and a non-killer strain *(Vibrio harveyi*) (S3 Fig). Moreover, we compete a killing deficient (T6SS-) variant of the *V. cholerae* strain against other T6SS killer strains (same four other environmental isolates of *V. cholerae* and *Enterobacter cloacae* [53, 54]) to test whether debris accumulation can be independent of the diverse toxins used by different strains (S4 Fig). In every strain combination, we observe the signatures of killing inhibition due to dead cell debris, i.e. localized dead cell stain (PI) that aligns with both the interface between strains as well as dark outlines in bright-field microscopy.

The evidence presented in Fig. 1 and Fig. 2 suggests that accumulated dead cell debris prevents contact between cells and thus prevents contact killing. However, these data cannot rule out counter-hypotheses that T6SS gene expression changes in response to cell density, altered nutrient density, or a change in oxygen concentration. To directly test if the presence of dead cell debris hinders killing, we mechanically disturb the structural organization of the biofilms and thus break down dead cell debris barriers, without otherwise altering biofilm conditions (see sketch in Fig. 3a). First, we study a co-culture of killer and target variants of *V. cholerae* (S1 Video). We inoculate the strains (OD_600_ = 1) between an agar plate and a cover glass. Killer cells initially expand their population, and then expansion halts as killing ceases (Fig. 3). After 4.5 hours we shear the biofilm by rotating the cover slip with respect to the agar pad in small circular motions (diameter ∼2mm), until both strains and the dead cell debris are well mixed. After the perturbation, we observe an immediate increase in the fraction of killer cells indicating that killing resumes (Fig. 3b). However, after the perturbation the remaining target cell domains are too small for us to observe if dead cell debris barriers again separate killer and target cells.

**Fig 3.**
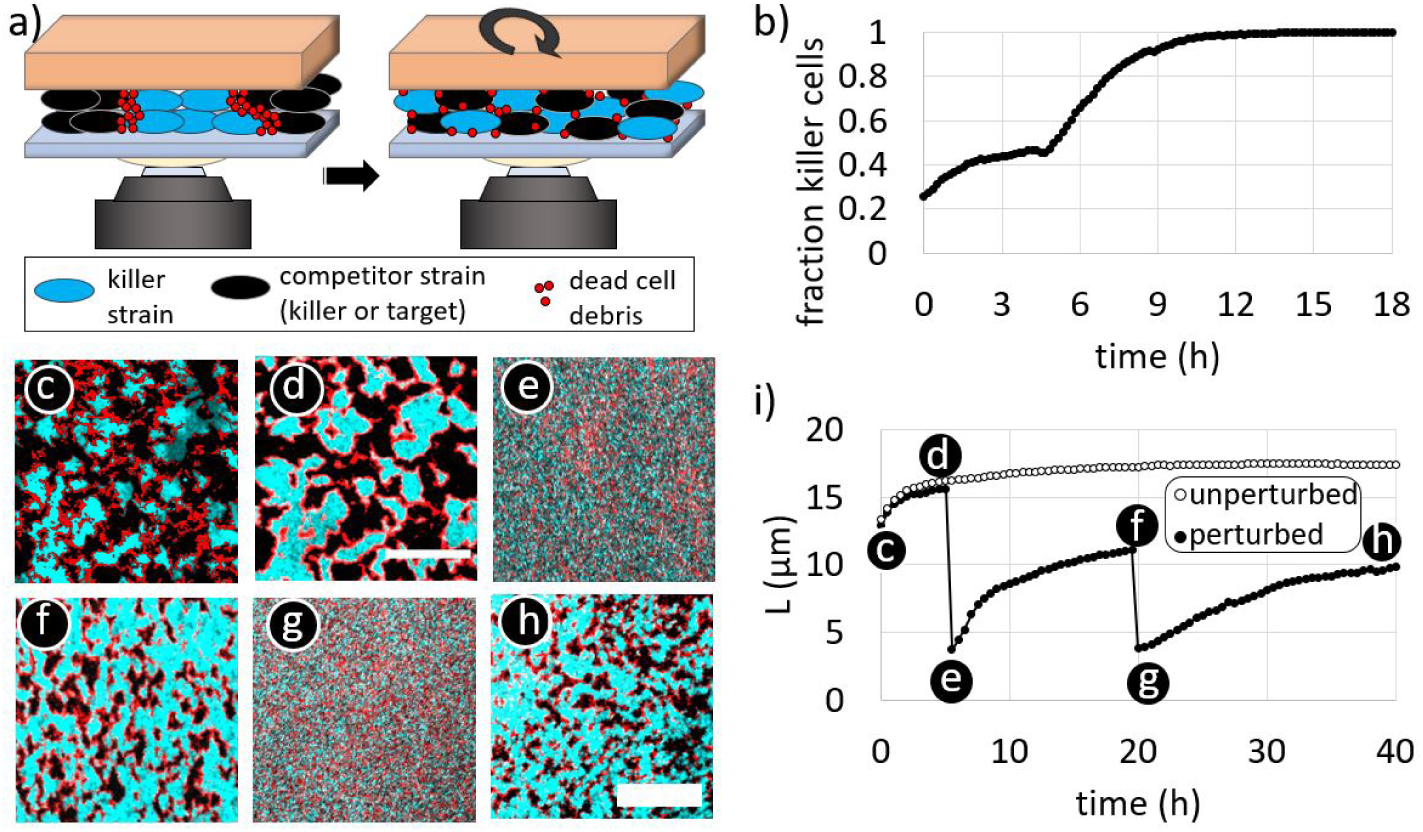
Mechanical perturbation of dead cell debris barriers between domains of competing strains. (a) Sketch of mechanical perturbation experiments. The agar pad is rotated, mixing both strains and the dead cell debris within the biofilm. (b) For unidirectional killing (*V. cholerae* killer against target strain), the fraction of killer cells saturates after an initial increase. After mechanical perturbation at 4h, the fraction of killers increases again as killing resumes, almost eliminating target cells (see also S1 Video). (c-h) Time-lapse fluorescence images of mechanical perturbation of a mutual killer biofilm show that phase separation occurs repeatedly: (c) after a cellular monolayer has formed (d) the biofilm after ∼5 hours of growth, (e) the biofilm immediately after the first mechanical perturbation, (f) the biofilm ∼15 hours after the first perturbation, (g) the biofilm immediately after the second perturbation, and (h) the final biofilm structure, 20 hours after the second perturbation. In the fluorescence images, one killer strain expresses sfGFP (cyan) while the other killer strain is unlabeled (appears black). In every image, the field of view is densely packed with cells. Dead cell debris is labeled with PI (red). (i) The mean domain size, *L*, in mutual killer biofilms increases with time, and suddenly decreases after mechanical perturbations at 5h and 20h. However, *L* increases after each perturbation, indicating that killing resumes (filled circles). In contrast, in un-perturbed biofilms *L* plateaus (empty circles).

To ensure that the domain size of each strain will remain large, we perform a new set of experiments competing two ‘mutual’ killer *V. cholerae* strains, i.e., each strain is T6SS+ and each strain is able to kill the other strain. We inoculate and grow a biofilm of mutual killers (Fig. 3c). After contact killing has nearly ceased (Fig. 3d), we shear the biofilm (Fig. 3e), thoroughly mixing the two strains and the dead cell debris. After mixing, we observe that large clonal domains again form over time, indicating that killing has begun again; further, we observe that these domains are separated by dead cell debris (Fig. 3f), and eventually killing again ceases. After shearing and mixing the biofilm a second time (at ∼ 20 hours, Fig. 3g), we again observe the growth of clonal domains that become separated by dead cell debris (Fig. 3h).

We quantify the growth, and mechanical destruction, of clonal domains by measuring the average size *L* of the domains of the fluorescent killer strain (Fig. 3i, filled circles) (see methods). *L* grows rapidly during the first ∼ 3 hours, and much more slowly after that time; upon mixing at 5 hours, *L* immediately drops to the size of about three cells (Fig. 3e,g). After that, *L* increases again, demonstrating that killing has resumed. We obtain a qualitatively similar trend after mixing the biofilm a second time, as indicated by an immediate decrease in *L*, followed by an increase. As a control, we measure *L* for a biofilm that is not mechanically perturbed. The mean domain size that emerges after ∼ 3 hours in the undisturbed biofilm increases by less than 8% over the next 37 hours (empty circles in Fig. 3h); in other words, the mean domain size is nearly constant after initial domain formation has occurred.

The above results show that the accumulation of dead cell debris can limit the utility of T6SS-mediated killing within biofilms. Contact killing initially eliminates opponent cells, structuring the biofilm population—but only until dead cell debris accumulates and killing nearly ceases. These observations suggest that the T6SS may have limited ability to facilitate biofilm invasion and that completely taking over a biofilm from a small number of T6SS+ cells would be unlikely. To test the ability of the T6SS to facilitate invasion, we examine the performance of single killer cells completely surrounded by target cells.

To do so, we inoculate and confine a dense cellular monolayer (OD_600_ = 10) of which 1% of the population are fluorescent killer cells. The remaining 99% of the cells are non-fluorescent, susceptible target cells, which are otherwise isogenic to the killers. We also perform control experiments in which the 1% fluorescent cells are defective killer cells (T6SS-). We find that after 24h of growth the final killer population is only ∼ 1.5X larger than the final fluorescent defective killer control population (Fig. 4b and c). In particular, clonal domains of killer cells have a mean size of 18.9*µm*^2^ (standard deviation of 26.7*µm*^2^); defective killer control cells form clonal domains with a mean size of 10.6*µm*^2^ (standard deviation of 14.9*µm*^2^). Thus, while killer cells expand their population more than killing-deficient cells, they are incapable of invading the existing biofilm and eliminating their competitors.

**Fig 4.**
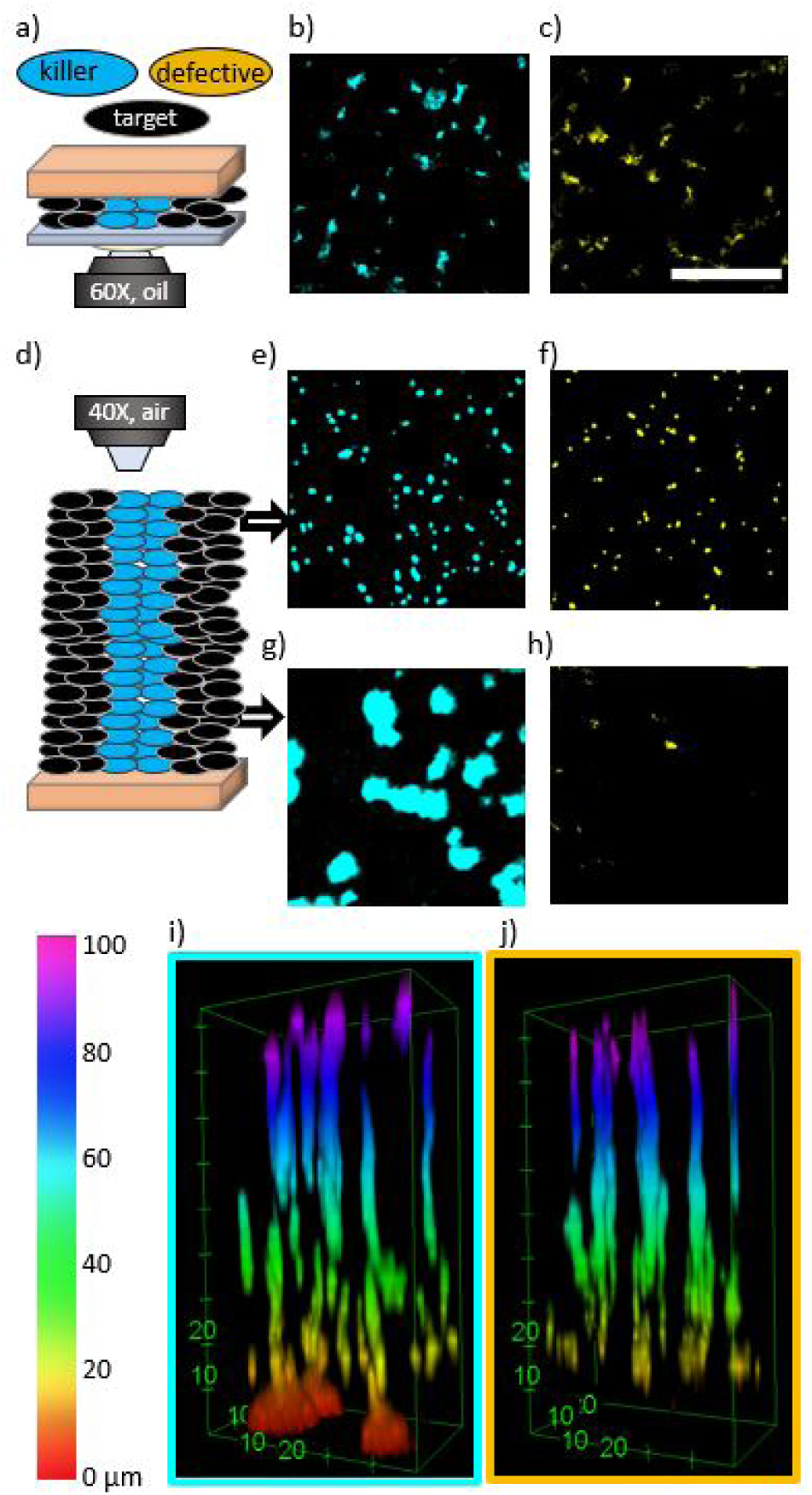
Invasion of dense biofilms by individual cells, shown by confocal microscopy images of sfGFP signal after 24 hours of biofilm growth. Biofilms grow from a dense cell monolayer containing 1% of cells that express sfGFP. These fluorescent cells are either T6SS+ killer cells (marked blue) or T6SS- defective killer cells (marked yellow) in control experiments. The remaining 99% of the population are non-fluorescent target cells, appearing black. In a confined geometry (see schematic in a), killer cells (b) only expand their population ∼1.5 times more than defective killer cells (c). For unconfined biofilms (schematic in d) the final abundance of fluorescent cells depends on distance above the agar surface. At the top of the biofilm, killer cells (e) only expand their population 1.1 times more than defective killer cells (f). At the bottom of the biofilm, killer cells (g) expand their population 275 times more than defective killer cells (h), which are almost absent. (Scale bar b,c,e-h: 50*µ*m) (i,j) Full 3D biofilm stack of fluorescent cells showing that killer (i) and defective killer (j) cells perform substantially different only within the bottom ∼10*µ*m. The distance from the agar pad is color-coded, see color bar.

Up to this point, all experiments are performed in confinement, i.e., biofilms are grown confined between an agar pad and a glass cover slip, to optimize the set-up for microscopy and exclude height-dependent differences [37, 55]. While bacteria often inhabit confined geometries in natural settings [56], it is unclear if confinement itself impacts population dynamics. Thus, we now explore contact killing in unconfined environments, by growing biofilms without a cover slip limiting their height. We again observe that the killer population rapidly increases for the first 3 hours, at which point killing slows substantially (S1 Fig c,d) and a thick layer of dead cell debris separates competing strains. To examine the ability of the T6SS to facilitate invasion of unconfined biofilms, we again inoculate a dense monolayer (OD_600_ = 10 inoculant) containing 1% fluorescent killer (T6SS+) or defective killer (T6SS-) cells and 99% unlabeled target cells. For both killer and defective killer experiments, biofilms reach a height of 92*µ*m*±*2*µ*m after 24 hours of growth. Similar to the results under confinement, the relative abundances of killer and non-killer control populations differ only slightly (Fig. 4 b,c); killer strains outperform non-killer strains by a factor of only 1.64 throughout the whole biofilm.

However, we observe that killer and defective killer strains exhibit markedly different behavior near the agar surface. At the top of the unconfined biofilm, the killer population is only 1.16 times greater than the defective killer population. The mean size of clonal domains are 8.0*µm*^2^ (standard deviation 6.0*µm*^2^) and 7.4*µm*^2^ (standard deviation 4.9*µm*^2^) for killer cells and defective killer cells, respectively. At the bottom of the biofilm (next to agar surface), the killer cells have expanded their population substantially; the mean diameter of clonal domains of killer cells within 4*µ*m of the agar surface is 115.4*µm*^2^ (standard deviation 146.7*µm*^2^). In contrast, the defective killer cells are nearly eliminated at the bottom (Fig. 4h). (Importantly, the remaining space in Fig. 4e-h is occupied by a non-killer strain lacking fluorescent proteins.) In the representative biofilm stacks shown in Fig. 4i and j, within 4*µ*m of the agar surface, the final population of killer cells is 270 times greater than the population of defective killer cells. Thus, although killer cells are only slightly better at invading unconfined biofilms than defective killer cells, killer cells are able to capture territory at the inoculation surface more efficiently.

## Discussion

Contact killing is common in bacterial biofilms [4]. While the lethal activity of T6SS is well known [7, 19, 31, 57, 58], T6SS-proficient bacteria rarely completely eliminate target cells – even if the killer strain starts with more cells than the target strain and experiments are run for long times [23]. We find that the efficacy of contact-killing in biofilms is limited by its material consequences. We show that killing-induced changes in the cell populations slow after approximately three hours, independent of the initial or final relative abundances of killer cells.

A reduction in the rate of T6SS killing is expected to some extent as T6SS killing only occurs when opposing strains are in direct physical contact. As target cells die and clonal domains grow in size, the amount of contact between opposing strains decreases. However, we find that expansion of the killer population slows dramatically after approximately three hours independently of the amount of contact between strains (which varies by a factor of ∼ 25 across the different cases explored here, Fig. 1). In fact, even when the T6SS-proficient strain has captured over 99% of the population, small domains of target cells, only a few microns in size, persist for hours (top curve in Fig. 1c). We confirm that the timing of slowed population changes coincides with the accumulation of dead cell debris between competing strains. By mechanically disrupting barriers of dead cell debris, we directly show that killer cells maintain their T6SS activity after population dynamics dramatically slow down. In these mechanical perturbation experiments, we only modify the positions of the two strains and the dead cell debris. Therefore, the accumulation of dead cell debris at the interface between strains is responsible for the cessation of T6SS killing.

The accumulation of dead cell debris thus has important implications for contact killing competitions. It was previously thought that T6SS mediated contact killing primarily facilitates bacterial competition and may “virtually eliminate” [58] susceptible competitor cells of similar number within a few hours [9, 29, 31, 58, 59] (unless target cells reproduce sufficiently fast [26]). In contrast, we observe that T6SS-mediated killing can instead facilitate the *coexistence* of antagonistic strains. Once formed, the dead cell debris barrier stabilizes the interface between opposing strains. This phenomenon is starkest in populations that combine killer cells and non-killer target cells. The target strain cannot fight back, but the accumulation of dead cell debris eventually prevents killing and protects the target strain from elimination. Depending on the initial killer fraction, the target population may only decrease by a few percent, even in biofilms that persist over long times.

At first sight, findings presented here might appear to disagree with conclusions drawn from T6SS competition killing assays, in which T6SS killing decreases the abundance of target cells by several orders of magnitude [7, 13, 15, 29, 31]. However, killing competition assays often start with a majority of killer cells over target cells, and the final abundance of target cells has a non-linear relationship with the initial abundance of killer cells. Decreasing the initial abundance of killer cells leads to enhanced survival of the target cells [23]. This non-linear phenomenon agrees with our microscopy results; we find that for the same pair of strains, target inhibition can be as high as 99.97% (three orders of magnitude) or as low as 79.47% (less than one order of magnitude) as the initial fraction of killer cells is varied from 0.50 to 0.09, respectively. In fact, despite the excess of killer cells in competition assays, the target strain is rarely, if ever, *completely* eliminated (if completely eliminated, CFU counts would be zero) [7, 13, 15, 29, 31]. Previous time-lapse competition assays found that the number of surviving target cells remains remarkably constant after a quick decline in the first few hours [23], consistent with the temporal dynamics we observe via microscopy. Therefore, our observations are in agreement with previously reported competition assays. Combined, these results demonstrate that the killing efficiency must be studied and discussed in context, with the initial abundances considered. For example, traditional competition assays show that, for some strains, a large number of killer cells can kill a small number of target cells very effectively, but they do not show how a small number of killer cells would perform against a large number of target cells.

It is important to note that the efficacy of T6SS killing can vary among different combinations of strains. In fact, environmental and clinical isolates of *V. cholerae* have been identified with very high and very low killing rates [29], suggesting that T6SS killing may be a highly effective killing mechanism in some but not all scenarios. Our findings suggest that the efficacy of T6SS killing could increase if cells can overcome the dead cell debris barrier. This could occur through a variety of processes. The barrier may be removed when dead cell debris is consumed by predators [60], broken down by secreted enzymes (such as lipases or DNAses), removed via shear flow [61], or dispersed by mechanical perturbations, among other potential mechanisms. The rate at which dead cell debris breaks down may also be impacted by the action of the delivered toxins themselves [62], which are known to exhibit a wide range of effects, from growth inhibition to lysis [20, 21, 35, 63, 64]. Further, while we observe similar characteristics of cell debris accumulation across different co-culture competitions containing *V. cholerae, E. cloacae*, and *V. harveyi*, the material and physical characteristics of dead cell debris may vary across different combinations of competing strains or species, impacting the stability of the barrier. Finally, motile cells could potentially penetrate barriers via migration. Thus, if mechanisms for rapidly breaking down or penetrating dead cell debris barriers are present, it remains possible that the T6SS could facilitate invasion of established communities or elimination of competitors.

Even though its efficacy is limited by the accumulation of dead cell debris, the utility of the T6SS remains quite high. First, contact killing initially facilitates the formation of clonal domains. Depending on the diffusion length of secreted goods [65], this initial domain formation may be sufficient to favor intra-strain cooperation [24, 66, 67] or reciprocal benefits between different strains [2, 68]. Second, contact killing can prevent social cheaters from emerging in a population. Such ‘policing’ effects have been observed in cases where social behavior is linked with T6SS regulation [69]. Finally, we find that contact-killing strains are able to capture and maintain territory near the surface on which they were inoculated; T6SS-deficient cells have a much lower probability of remaining near the surface.

The relative absence of killing-deficient cells from the nutrient dense surface results from mechanical interactions in biofilms. As cells reproduce they mechanically exert forces on neighboring cells [70] (similar to the interaction with dead cell debris in Fig. 2g). In dense monolayers, cells tend to migrate ‘up,’ as lateral motion is strongly limited by surrounding cells [70–73]. Once a cell is pushed up, it is statistically less likely to migrate ‘down.’ Hence, only a small number of defective killer cells remain at the bottom during early growth. In contrast, killer cells can initially contact and kill their neighbors. This opens free space on the agar surface at early times of growth, allowing killer cells to expand their territory near the nutrient-rich agar substrate. Occupying nutrient rich territories may have long term benefits, beyond the scope of our experiments [74].

While we only perform experiments with T6SS killing, results presented here potentially apply to many other contact killing mechanisms [4]. In fact, comparable phenomena of inhibited killing were seen in contact dependent growth inhibition experiments, in which live but non-reproducing cells were observed to prevent further contact between antagonistic cells and their targets [75]. Further, similar phenomena may impact killing mechanisms that act over longer distances, e.g., via diffusible deadly bio-molecules [76], phages [77] and bacteriocins [78]. These killing mechanisms will typically only be effective within some diffusion length. Killing may thus be hindered if a large dead cell barrier–one longer than the diffusion length–forms. However, more experiments are necessary to establish the generality of the effects observed here.

It is striking that contact killing indirectly facilitates the coexistence of antagonistic strains [79]. These results suggest that the T6SS, and perhaps other contact killing mechanisms, may not prompt a microbial ‘arms race,’ but instead may stabilize diverse communities. Further, these results align with recent works which question the ecological purpose of bacterial production of antibiotics [74, 80]. Nevertheless, the fact that contact killing facilitates coexistence suggests that the impact of the T6SS in bacterial consortia is complex, and the T6SS is more than simply a potent weapon.

## Materials and methods

### Bacterial strains

For experiments presented in figures in the main text, we used the *qstR*^∗^ constitutive killer *V. cholerae* C6706 and derivatives of it (Tab 1). Isogenic mutual killer experiments competed SN441 and JT1052; JT1052 was engineered from C6706 by replacing the aux1 and aux2 clusters with clusters from an environmental *V. cholerae* isolate 692-79 [81]. Experiments with unidirectional killing employed a T6SS-active strain (T6SS+) SN441 against a target strain (JT1053), which was a T6SS-defective mutant (T6SS-) of JT1052, prepared via *vasK* deletion. The SN441 killer strain constitutively expresses a superfolder green fluorescent protein (sfGFP).

**Table 1.**
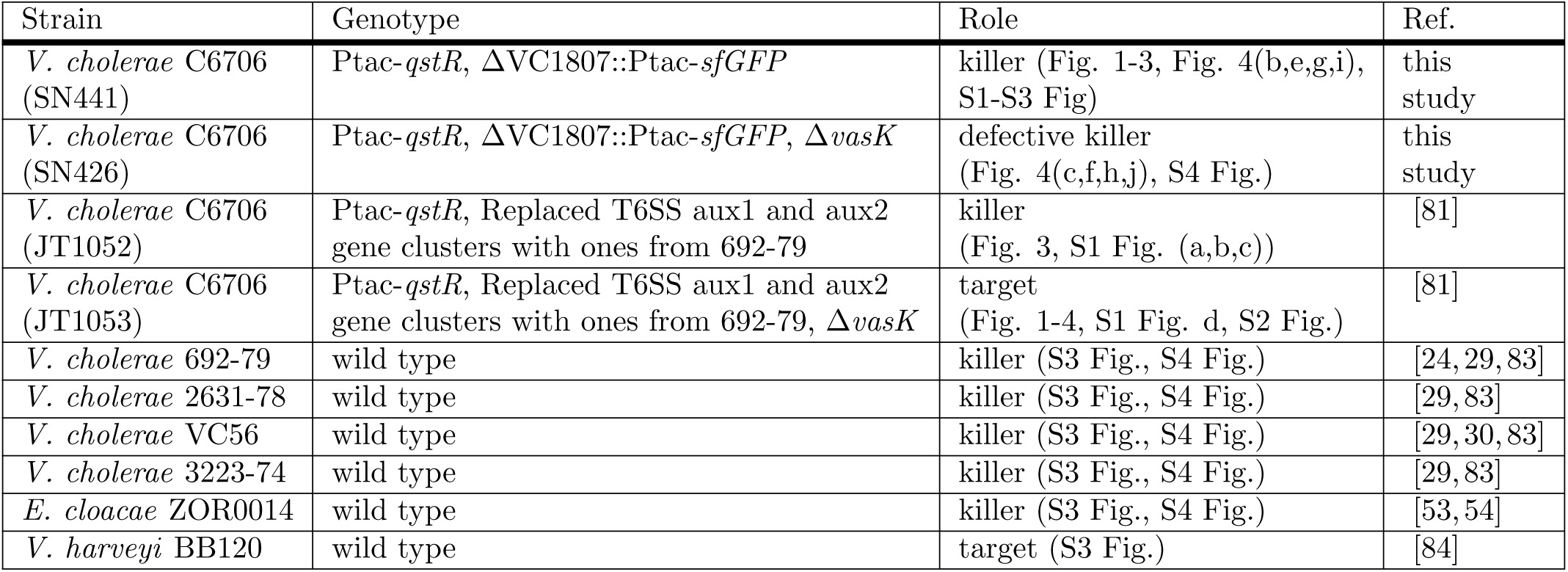
Strain names and specifications.

### Construction of genetically altered *V. cholerae* strains

All *V. cholerae* mutant strains were constructed using allelic exchange methods as described previously [82] and confirmed by PCR and sequencing. Restriction enzymes, polymerases and Gibson assembly mix reagents were used to manipulate DNA fragments and plasmids according to manufacturer instructions (New England Biolabs and Promega).

### Sample preparation

All bacterial strains (apart from *V. harveyi*) were grown overnight in lysogeny broth (LB), and then back-diluted in fresh LB for an additional 3h at 37°C. Only *V. harveyi* was grown overnight at 30°C in marine (LM) medium. Then, for each experiment liquid co-cultures of two strains were mixed, accordingly. For measurements of spatio-temporal dynamics in unidirectional killing (Fig.1c), we employed four different killer to target volume mixing ratios: 1:1, 1:2, 1:5, and 1:10. (Note that in Fig. 1, these ratios do not reflect the fraction of killer cells at 0 hours. While the samples in Fig.1 were inoculated at OD_600_ = 1, strain fractions were extracted only after a dense monolayer had formed, which then contains altered strain fractions.) For volume mixing ratios in all other experiments, refer to details given in the main text. Liquid cultures were either aliquoted OD_600_ = 1 or OD_600_ = 10 (see main text for each experiment). To achieve an effective OD_600_ = 10, cultures with an OD_600_ = 1 were centrifuged and re-suspended 1/10 the volume of fresh medium. After preparation of liquid co-cultures, they were inoculated on agar pads. We prepared 1.5% LB agar pads (1.5% LM agar pads used for *V. harveyi*) and added 7*µ*l of 100*µ*g/ml propidium iodide (PI) on the agar pad surface. After the PI droplet dried, 1*µ*l of liquid co-culture was inoculated on that same spot of the agar pad. For measurements in vertical confinement, a cover glass was placed on top of the agar pad directly after inoculation. All biofilms were grown under optimal conditions in either an Okolab caged incubator attached to the confocal microscope (time-lapse measurements) or a separate incubator box (single-time measurements after 24h of growth). Biofilms were grown at 37°C, but co-cultures containing *V. harveyi* were grown at 30°C.

### Microscopy

All images were recorded with a Nikon A1R confocal microscope. We used FITC and TRITC filters for recording the fluorescence signal from the sfGFP expressing cells and the PI-labeled DNA of dead cells. We used bright-field mode for recording bright-field images. The microscope is additionally equipped with an OkoLab incubator box (operated at 37°C and maximum humidity) to ensure optimal growth conditions. Samples in 2D confinement were imaged with a 60X oil immersion objective (NA=1.45) and a vertical step size of 0.5*µ*m. For unconfined growth, images were taken with a 40X long distance objective (NA=0.6) and with a vertical step size of 2*µ*m. For time-lapse recordings of the two-strain coarsening process, images were recorded every 10 minutes. Tracking of PI-labeled single dead cells was performed in 5 minutes sequences.

### Image processing and analysis

All image processing and analysis was performed with Fiji. The fraction of the fluorescent strain (Fig. 1c, Fig. 4) was extracted from threshold images: a threshold was applied to the the fluorescent image recorded from sfGFP signal, and the sum of pixels that contain sfGFP fluorescence intensity above the threshold was counted.

The mean relative PI intensity as a function of distance from the interface between strains (Fig. 2f) was determined as follows. Starting with the thresholded image of the sfGFP channel as described before, an automated edge detection was applied to identify the interface lines. After that, a mask was created that covers all area residing within a distance *r* from the interface lines. A separate mask was created for each distance analyzed. Each mask was applied to both the fluorescence image containing the PI signal and the bright-field image. We then measured the mean brightness intensity (PI or bright-field signal) within the area covered by each mask. This gives the mean brightness intensity for each distance. The obtained curve of mean brightness intensities as a function of distance was finally normalized to a range between 0 and 1. This is achieved by dividing each measured mean brightness by the mean brightness of the full image, subtracting −1, and then dividing by the brightness value at the smallest distance.

To perform measurements on individual dead cells, we detected single spots in images from the PI channel and measured the number of detected spots (Fig. 2h) and the mean size per spot (S2 Fig) per recorded time point.

The structural analysis of the phase-separation of two competing strains in Fig. 3 was performed via a Fourier transform analysis as described in [24]. In short, we applied a Fourier transformation on the fluorescence image of phase-separating strains. Using the transformed image, we calculated the structure factor *S*(*q*) by radial integration, determined the mean structure factor *S*_*m*_ and extracted the corresponding characteristic wave length *q*_*m*_ at *S*_*m*_(*q*_*m*_). Finally, the characteristic domain length was obtained as 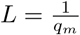.

For 3D image stacks of biofilms grown without confinement (Fig.4e-j), the fluorescence brightness was corrected with depth since the fluorescence intensity decreases with biofilm depth. Deeper biofilm layers experience less fluorescence excitation and emitted light undergoes diffraction which decreases detectable intensity. Cells deep in the biofilm may also express fluorescent proteins at a lower rate than cells near the surface. To characterize how the intensity of the fluorescent signal varies as a function of height, a full z-stack of a single-strain biofilm (containing only one fluorescent strain) was recorded. We measured the mean fluorescence intensity as a function of biofilm depth. Due to the huge intensity variation across a biofilm of 95*µ*m thickness, we recorded the biofilm stack in two sections with different settings of the microscopy laser; high laser power and gain were applied to image the lower section, and low values were applied for intermediate and upper sections. Using the same settings, we measured the co-culture biofilms; afterwards, we corrected the depth-dependent intensity according to the depth dependent intensity measured from the single-strain biofilm.

## Supporting information

**S1 Fig.**
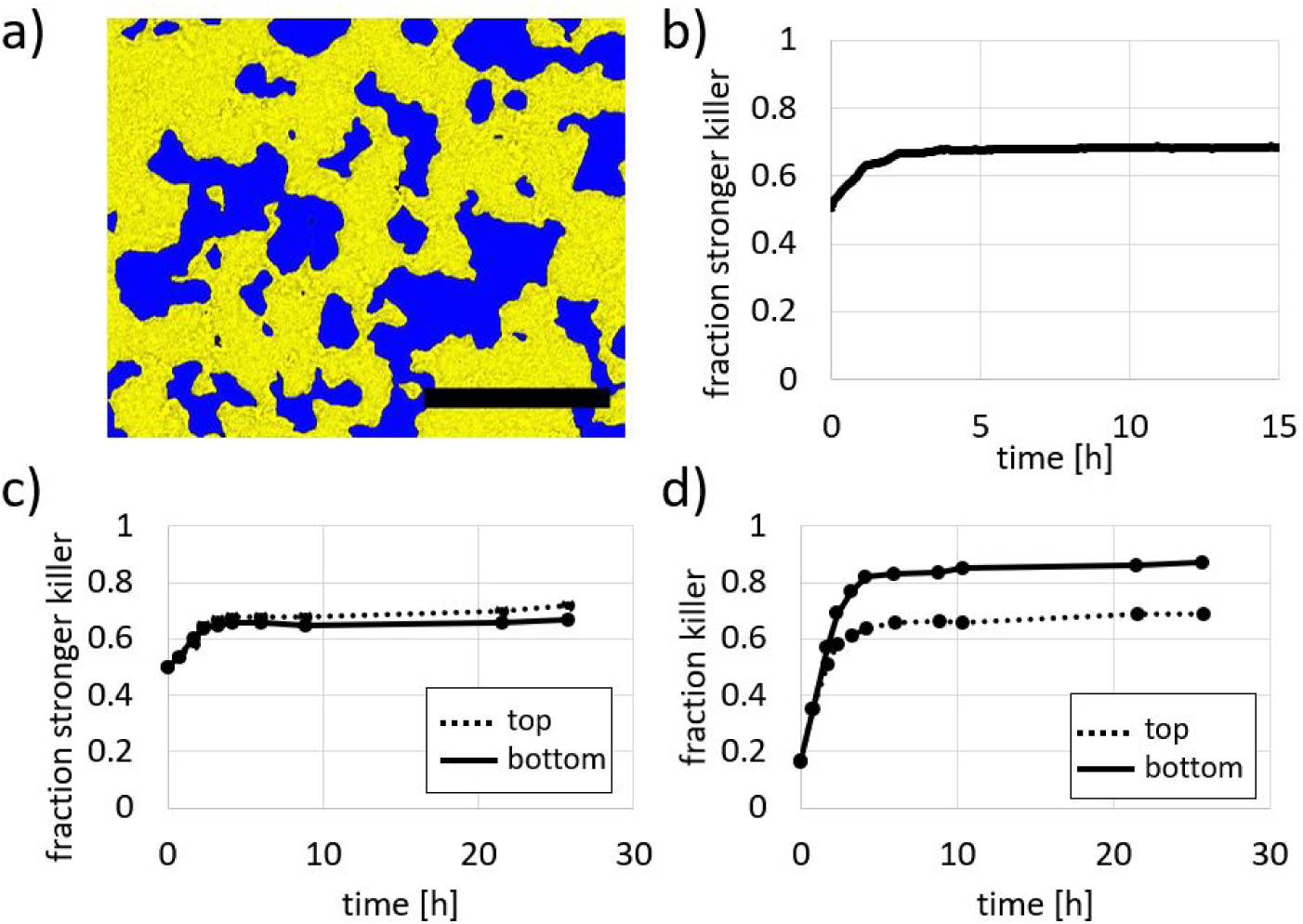
Spatio-temporal analysis of phase separation with and without confinement. a) T6SS-mediated contact killing between mutual killers (initial number ratio of 1:1) leads to the formation of clonal patches (S2 Video) as shown here by a fluorescent microscopy overlaid on a bright field image image. The observed domain formation agrees with previous findings [24]. The image is recorded after 7 hours of biofilm growth in 2D confinement. (Scale bar: 50µm) The fraction of the more potent killer increases at early times followed by a significantly slower increase after 3 hours in both cases for 2D confinement (b) and without confinement, i.e. without cover glass on top (c), where the latter shows almost identical fractions at the top and the bottom of the biofilm. (d) In the case of unidirectional killing in unconfined growth, we obtain a similar temporal behavior, but here the killer expands more at the bottom of the biofilm than at the top. This is consistent with findings in Fig. 4i.

**S2 Fig.**
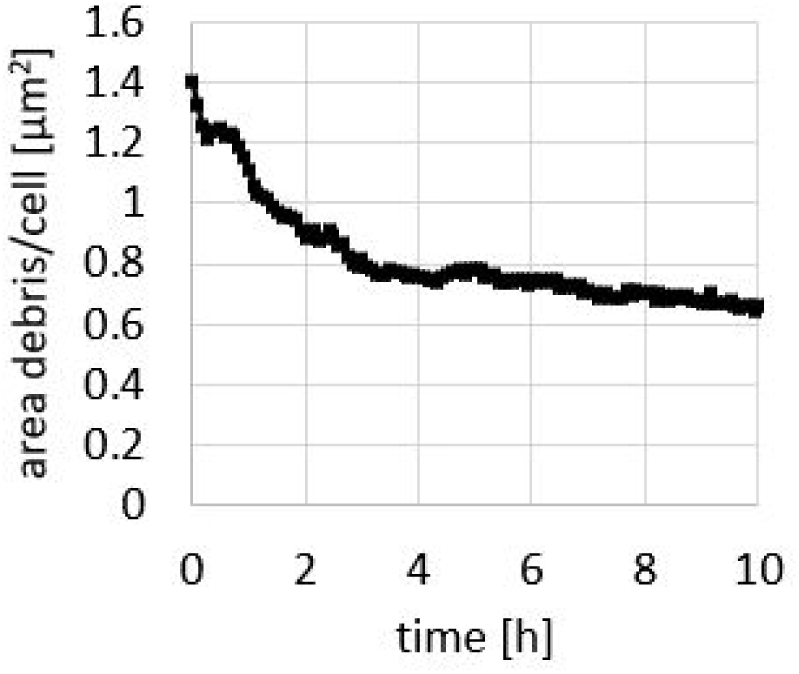
Persistence of dead cell debris. The PI-labeled area per dead cell is recorded over time by tracking individual dead cells over 10 hours and extracting the mean PI-labeled area per dead cell. After a quick decline (due to compression or degradation of cell debris), the amount of labeled area saturates, demonstrating that PI-labeled DNA persists for hours after cell death. The data are obtained from a dense mix of killer and target cells (number ratio of 50:1, inoculated at OD_600_ =10), ensuring that most dead target cells are sufficiently far from each other to enable long-time tracking.

**S3 Fig.**
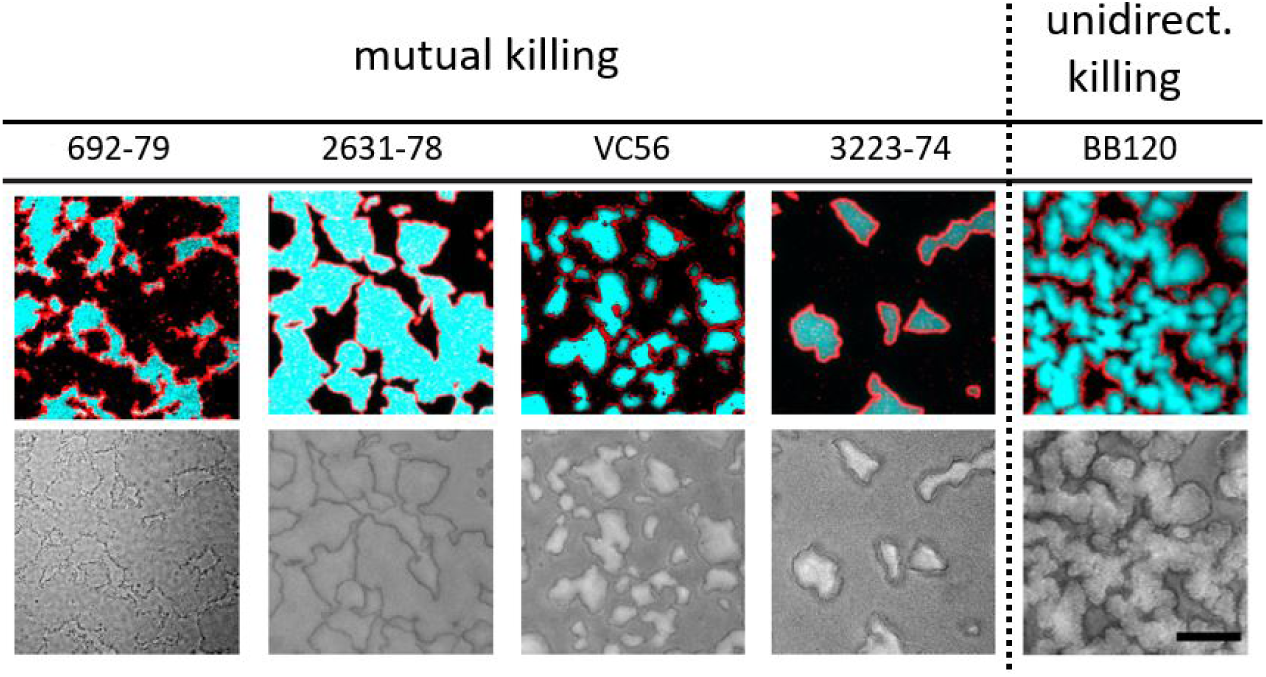
Competition outcomes from mutual and unidirectional killing between nonisogenic competitors, involving the C6706 T6SS+ killer strain. The C6706 T6SS+ killer strain is competed against other T6SS+ killers (mutual killing) and non-killer target strains (unidirectional killing). The top row depicts fluorescent microscopy images containing the fluorescent C6706 killer strain (cyan) and PI-labeled cell debris. The T6SS-proficient competitor strains (four different *V. cholerae* strains) and the non-killer competitor strain *V. harveyi* are not fluorescent and appear black. The bottom row depicts the same areas of view but as brightfield images. For all strain combinations, we observe that PI-labeled dead cell debris accumulates at the interface between competitors, and optical intensity variations occur at the interface between strains in the bright-field images (though please note, non-isogenic strains exhibit different intensities in bright-field imaging, partially, but not completely, obscuring the dark outlines between strains). For each strain combination, the initial number ratio between C6706 and its competitor strain was adjusted (ranging from 1:1 to 1:5) such that large domains of each strain were observed after 24 hours of biofilm growth. All co-cultures were incubated at OD_600_ = 1. Scale bar: 50*µ*m.

**S4 Fig.**
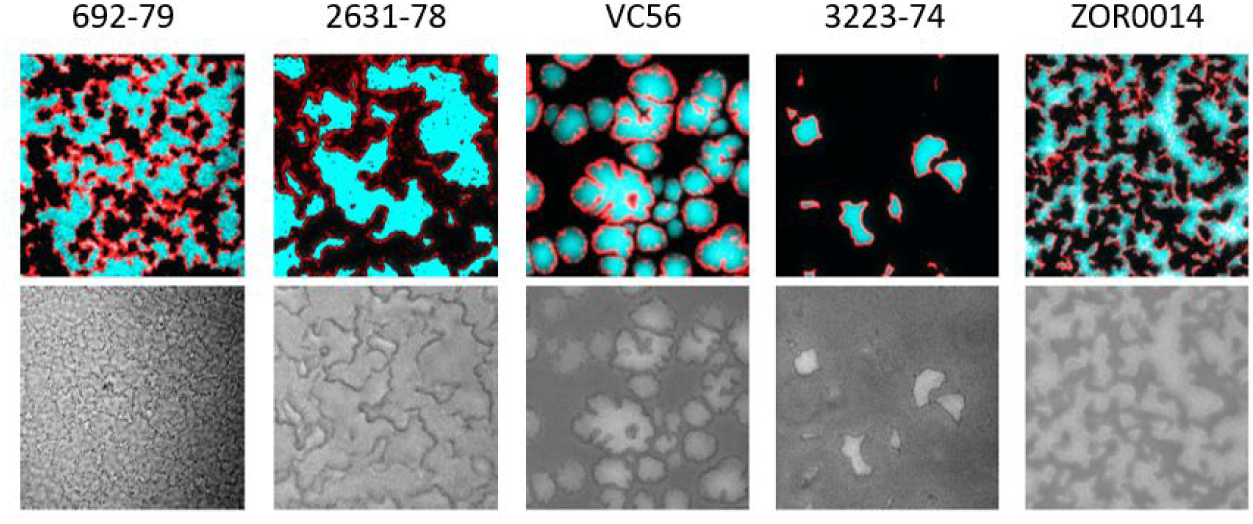
Unidirectional competition outcomes between a T6SS-defective C6706 strain and other, non-isogenic T6SS+ killer strains. The top row depicts fluorescent microscopy images containing the fluorescent C6706 killer strain (cyan) and PI-labeled cell debris. The T6SS-proficient competitor strains (four different *V. cholerae* strains and *E. cloacae* (data not published)) are not fluorescent and appear black. The bottom row depicts the same areas of view but as brightfield image. For all strain combinations, we observe that PI-labeled dead cell debris accumulates at the interface between competitors, and optical intensity variations occur at the interface between strains in the bright-field images. Please note, non-isogenic strains exhibit different intensities in bright-field imaging, partially, but not completely, obscuring the dark outlines between strains. This shows that the accumulation of dead cell debris is a general feature of T6SS mediated killing and occurs from a variety of toxins. For each image the initial number ratio between C6706 and its competitor strain was adjusted (ranging from 1:3 to 1:5) such that large domains of each strain were observed after 24 hours of biofilm growth. All co-cultures were incubated at OD_600_ = 1.

**S1 Video. Mechanical perturbation in a biofilm with unidirectional killing.** Time-lapse images of a vertically confined, growing *V. cholerae* population composed of a T6SS+ killer strain and a T6SS-target strain recorded with confocal microscopy. The recordings show brightfield images with overlaid fluorescence signal from the killer strain (cyan); the target strain is not fluorescent. The time, given in hours, gives negative values for the period before a dense biofilm has formed. Positive values count the time after dense packing of cells has been obtained.

**S2 Video. Population dynamics in a biofilm composed of mutually killing *V. cholerae* strains.** Time-lapse images of a vertically confined biofilm composed of T6SS-proficient, mutually killing *V. cholerae* strains recorded with confocal microscopy. The recordings show brightfield images with overlaid fluorescence signal from one killer strain (cyan); the other killer strain is not fluorescent.

**S3 Video. Accumulation of dead cell debris in a biofilm composed of mutually killing *V. cholerae* strains.** Time-lapse images of a vertically confined biofilm composed of T6SS-proficient, mutually killing *V. cholerae* strains recorded with confocal microscopy. The recordings show brightfield images with overlaid fluorescence signal from one killer strain (cyan); the other killer strain is not fluorescent. Dead cell debris appears red from PI labeling.

## Acknowledgments

GS is grateful for funding from the German National Academy of Sciences Leopoldina. BKH acknowledges funding from the Gordon and Betty Moore Foundation (Grant No. 6790.13), the School of Biological Sciences Abell Fellowship, and the College of Sciences Cullen Peck Scholars Award. PJY acknowledges funding from the Coulter Foundation and the Georgia CTSA.

